# Consciousness is indexed by analogous cortical reorganization during sleep and anesthesia

**DOI:** 10.1101/2022.11.15.516653

**Authors:** Bryan M. Krause, Declan I. Campbell, Christopher K. Kovach, Rashmi N. Mueller, Hiroto Kawasaki, Kirill V. Nourski, Matthew I. Banks

## Abstract

Although sleep and anesthesia are predicted to share common neural signatures of transitions into and out of unconsciousness, supportive evidence has been elusive. We identified these signatures using intracranial electroencephalography in neurosurgical patients. We applied diffusion map embedding to map cortical location into a space where proximity indicates functional similarity using a normalized connectivity (‘diffusion’) matrix, itself a rich source of information about network properties. During reduced consciousness, diffusion matrices exhibited decreased effective dimensionality, reflecting reduced network entropy. Furthermore, functional brain regions exhibited tighter clustering in embedding space with greater distances between regions, corresponding to decreased differentiation and functional integration. These changes were not region-specific, suggesting global network reorganization. These results strongly suggest common neural substrates for loss and recovery of consciousness during anesthesia and sleep, providing a systems-level mechanistic understanding within an intuitive geometric context and laying the foundation for evaluation of cortical state transitions in clinical settings.

## Introduction

Leading theories of brain function predict that loss and recovery of consciousness (LOC, ROC) are precipitated by large-scale reorganization of cortical networks. This reorganization might result in altered functional integration and differentiation in the brain^1^, communication into or out of prefrontal cortex and amplification of sensory signals^2^, or feedback connectivity^3^ with concomitant effects on predictive processing^4^.

It is postulated that changes in the brain underlying LOC and ROC should overlap regardless of the circumstances of their occurrence^5,6^, and thus comparison of changes in the brain in multiple contexts can be informative^7^. For example, general anesthesia and natural sleep exhibit common behavioral and physiological features, including both dreaming (i.e., conscious experience while disconnected from the environment) and unconsciousness^8–10^ and decreased cerebral blood flow and metabolic rate^11–13^. Recently, we observed similar changes in cortical functional connectivity during anesthesia and sleep. Stages of higher probability of consciousness, including wake, propofol sedation, and N1 and REM sleep, exhibited connectivity profiles that were similar to each other but distinct from stages of reduced probability of consciousness, including propofol unresponsiveness and N2 and N3 sleep^7^. These findings were consistent with a network transition boundary for consciousness common to anesthesia and natural sleep, which we linked to the ongoing debate about the locus of the neural correlates of consciousness^14,15^. However, the degree to which the brain traverses an overlapping complement of network states during anesthesia versus sleep remains controversial^16,17^.

Questions remain as well about the specific features of network reorganization associated with transitions into and out of consciousness. For example, although altered network connectivity is observed consistently during both anesthesia and sleep, some studies report that connectivity is decreased^18–24^ and others that it is increased^25–27^. Furthermore, selective effects of anesthesia have been reported on both feedback^19,28^ and feedforward^29^ connectivity. Decreased thalamo-cortical connectivity has been reported for both anesthesia and sleep^30,31^, but at least for anesthesia this change is unlikely to be causal for LOC^11^. Distinct effects of anesthesia and sleep also have been reported in analyses of resting state networks (RSNs). Increased modularity of RSNs during NREM sleep was reported to be accompanied by greater connectivity overall^27^, suggesting differential effects on within-versus between-network connectivity. By contrast, during propofol anesthesia both between- and within-network connectivity was observed to decrease^32,33^.

The organizational features of the conscious brain can be couched in terms of the balance between integration and differentiation, i.e., between the unified nature of conscious experience and its vast potential for variation. Investigations that operationalize changes in integration and differentiation during anesthesia and sleep have produced consistent results. Decreased differentiation of brain activity has been reported, indexed by reduced complexity of both evoked brain responses^34–37^ and of spontaneous brain signals^38,39^. Brain functional integration decreases during both anesthesia^40^ and NREM sleep^27^; surprisingly, however, overall connectivity decreased in the former study and increased in the latter.

In a small number of studies, data recorded during anesthesia and sleep has been compared directly^17,25,41^, but no clear and consistent mechanism for loss of consciousness emerges from these analyses. We seek a unifying framework for understanding network reorganization during the different stages of anesthesia and sleep and relating these changes to theoretical constructs. Here, we explore the functional geometry of cortical networks using diffusion map embedding (DME)^42^ and show that changes in connectivity across stages of sleep and anesthesia reflect changes in the organization of cortical networks that may contribute to loss of consciousness. As in our previous work, we distinguished stages corresponding to substantially reduced probability of consciousness (propofol unresponsiveness, NREM sleep), from the wake state and from stages of higher probability of conscious experience (propofol sedation, light sleep, REM sleep). We show that entry into states of reduced consciousness during both anesthesia and sleep can be indexed reliably by a single parameter, the effective dimensionality of the normalized connectivity matrix. We present an analytical framework that provides an intuitive, geometric understanding of changes in cortical networks associated with states of reduced consciousness and observed reductions in effective dimensionality. Globally, brain regions become more distinct (reduced *functional integration*), moving farther apart in functional embedding space. Locally, brain subregions become less distinguishable (reduced *differentiation*), moving closer to each other in the functional embedding space. This unifying framework has a practical utility in identifying cortical state transitions in clinical settings and broader implications for understanding the neural basis of consciousness.

## Results

### Summary of experiments and recordings

Resting state iEEG recordings were obtained in neurosurgical patients undergoing intracranial monitoring for the purpose of identifying seizure foci. Demographic information and electrode coverage are summarized in **Supplementary Tables 1** and **2**. Typical electrode coverage is shown for one participant **in Figure 1a**, and a summary of the brain parcellation scheme and of electrode coverage across all participants are shown in **Supplementary Figure 1**. Each recording site was assigned to a region of interest (ROI), color-coded according to a functional parcellation scheme illustrated in **Supplementary Figure 1a**. This scheme was derived from ana analysis of daytime resting state iEEG data from a complementary dataset obtained during daytime wake^43^. To investigate changes in cortical network organization during transitions in arousal and awareness, data were recorded during induction of propofol anesthesia just prior to removal of electrodes (*N* = 14 participants; **Supplementary Figure 2**), and during overnight sleep (*N* = 15 participants; **Supplementary Figure 3**). As in our previous work, we identified stages of anesthesia (WA: pre-drug wake; S: sedated but responsive; U: unresponsive) using a standard clinical assessment tool (Observer’s Assessment of Arousal Score; OAA/S^7,44^). Sleep stages were identified using standard polysomnography (WS: wake; N1: light sleep; N2 and N3: NREM; R: REM).

**Figure 1.**
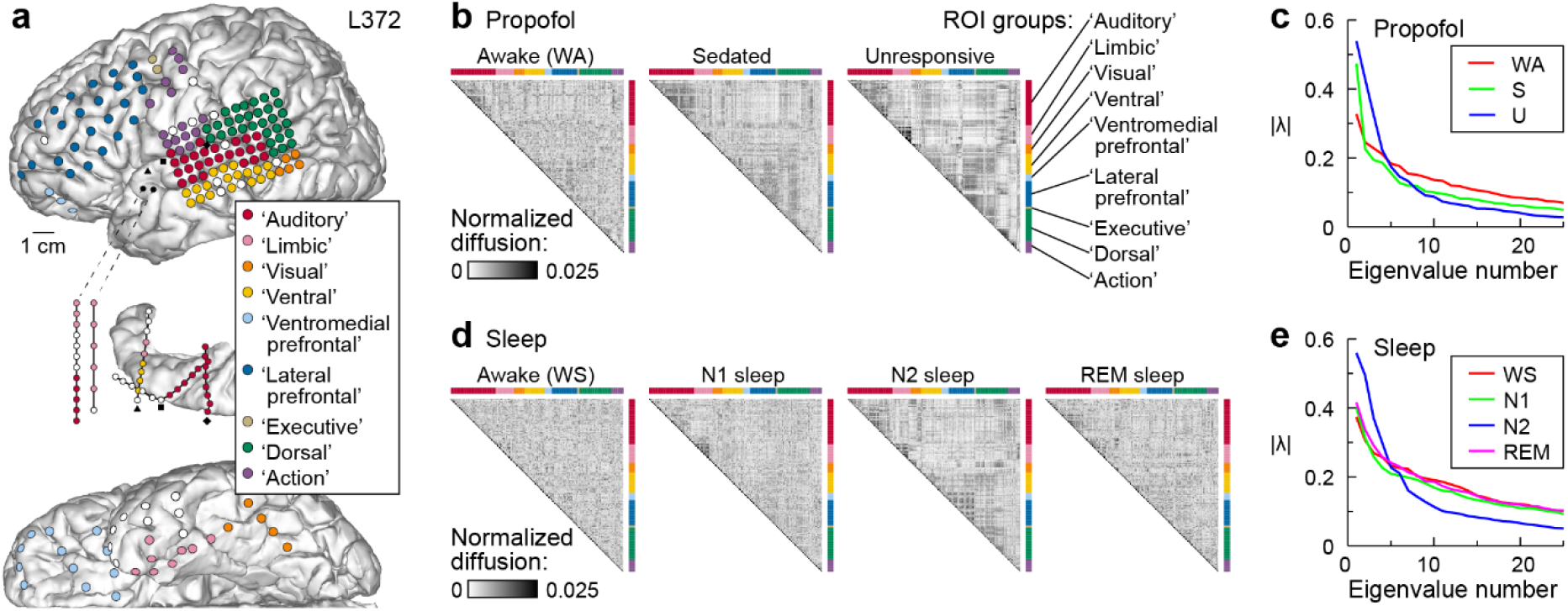
Network organization varies during anesthesia and sleep. **a**: Typical electrode coverage in one participant (L372). Recording sites are color-coded according to the ROI group. White symbols denote sites excluded from the analysis due to excessive noise, artifacts, location within seizure focus, in white matter, or outside the brain. Black symbols denote depth electrode insertion points. **b**: Diffusion matrices **P_symm_** during propofol anesthesia for the participant in **a**. Each matrix is from one minute of data. **C**. Spectra of **P_symm_** calculated from the example matrices in **b**. For these examples, *D*_E_(WA) = 0.30, *D*_E_(S) = 0.20, *D*_E_(U) = 0.10. **d**: Diffusion matrices **P_symm_** during sleep for the participant in **a**. Each matrix is from one minute of data. **e**: Spectra of **P_symm_** calculated from the matrices in **d**. For these examples, *D*_E_(WS) = 0.31, *D*_E_(N1) = 0.30, *D*_E_(N2) = 0.14, *D*_E_(REM) = 0.32. For panels **b**-**e**, data recorded during anesthesia and sleep experiments were divided into segments of length 60 s, and the diffusion matrix and spectrum computed for each segment. Matrices and spectra shown are from the segments with effective dimensionality closest to the median value for each stage of anesthesia and sleep in this participant.

### Changes in cortical network organization during anesthesia and sleep

Functional connectivity was calculated as orthogonalized gamma band power envelope correlations^43,45^, yielding for each one-minute data segment an electrode × electrode connectivity matrix. The first steps of DME analysis are to create a similarity matrix by applying cosine similarity to the functional connectivity matrix, then normalize, threshold, and make symmetric the similarity matrix to yield a diffusion matrix **P_symm_**. **P_symm_** describes the diffusion of an input signal applied to nodes (i.e., recording sites) on the graph^42^. When **P_symm_** is sorted by brain region (indicated by colored bars in **Figure 1b**), increasing community structure in the graph becomes evident in states of reduced consciousness under propofol anesthesia (sedated/S, unresponsive/U). The degree of community structure can be quantified by examining the eigenvalue spectrum of **P_symm_** (**Figure 1c**). Random graphs, i.e., those with maximal entropy, have spectra that are approximately flat. Graphs with strong community structure have spectra that are more peaked. The underlying entropy of the graph, and hence the shape of the spectrum, can be quantified using the effective dimensionality *D*_E_ ∈ (0,1), a function of the eigenvalue spectrum and a graph theoretic measure of complexity (see Methods). Importantly, the eigenvalue spectrum and calculation of *D*_E_ do not require nodes to be ordered or classified. Like anesthesia, NREM sleep was associated with a more structured **P_symm_** and more peaked spectrum (**Figure 1d,e**).

The time series of *D*_E_ computed for each 60-second data segment recorded during an experiment reveals striking changes in network structure over time during both anesthesia and sleep experiments, with transitions into S and U during anesthesia and into N2 during sleep accompanied by sharp decreases in *D*_E_ (**Figure 2**). Notably, *D*_E_ was consistently high during stages associated with higher probability of consciousness (WA, S, WS, N1, REM). Two additional examples from participants recorded during both propofol anesthesia and during sleep are shown in **Supplementary Figure 4**.

**Figure 2.**
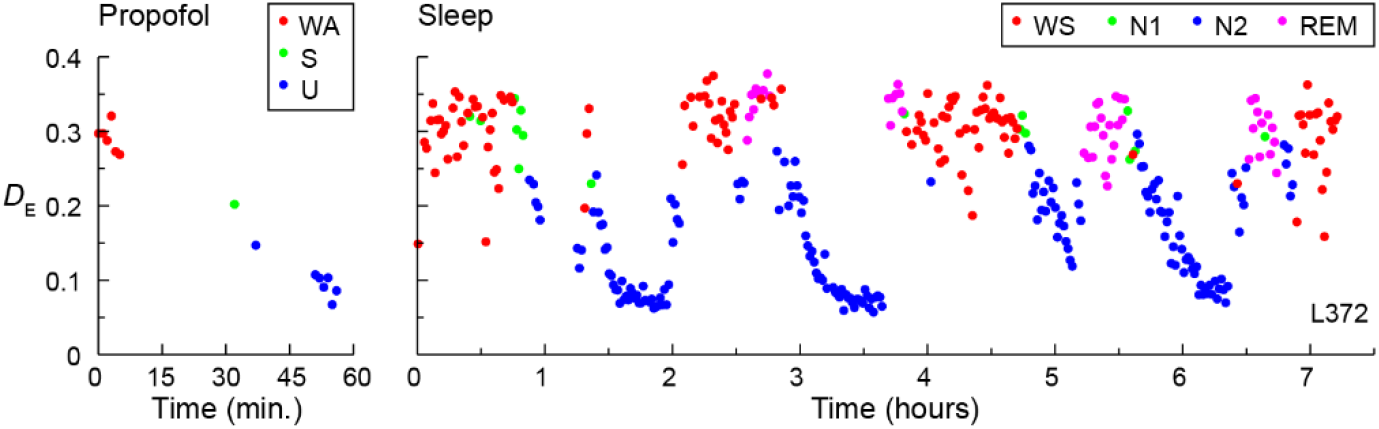
Time series of *D_E_* from an example participant. Changes during anesthesia and sleep are shown in left and right panel, respectively. Each data point represents one minute of data. Time is depicted relative to the start of recording. Same participant(L372) as in **Figure 1**.

Data were summarized across participants by first averaging *D*_E_ within participant across all segments associated with each stage of anesthesia and sleep. *D*_E_ varied significantly by state for both propofol anesthesia and sleep (likelihood ratio test for omitting state: propofol χ^2^(2) = 42.0, p < 0.0001; sleep χ^2^(4) = 79.3, *p* < 0.0001) (**Figure 3**). For propofol anesthesia, mean *D*_E_ decreased progressively from WA to S to U (**Supplementary Table 4**). During sleep, *D*_E_ for N1 and R were not significantly different from WS, but *D*_E_ decreased in N2 and decreased further in N3 (**Supplementary Table 4**). These results were robust to the choice of threshold in calculating **P_symm_** (**Supplementary Figure 5**).

**Figure 3.**
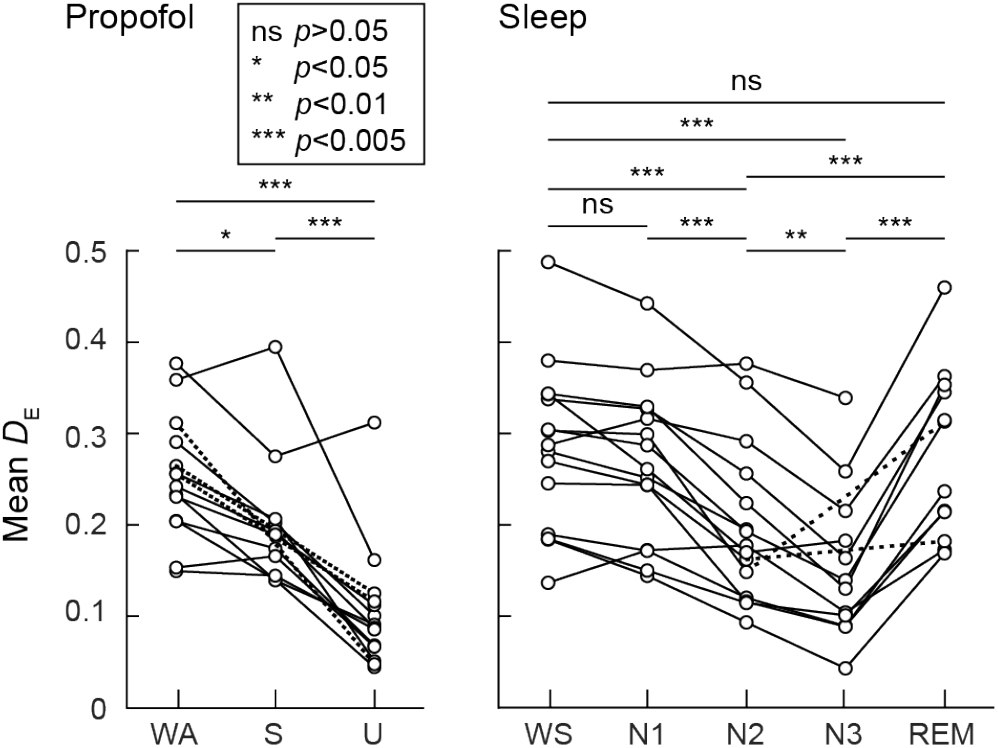
Summary of changes in *D*_E_ in states of reduced consciousness. Changes during propofol anesthesia and sleep are shown in left and right panel, respectively. Symbols are mean within a participant, connected by lines for data points from the same participant. Dashed lines are used to connect points when data in the intervening state (S; N3) is not available for that participant. *P*-values are from paired post-hoc comparisons, adjusted using multivariate *t*-distribution.

### Changes in cortical network organization are not regionally specific

It is plausible that observed changes in *D*_E_ between network states could be dominated by changes in a subset of recording sites in specific brain regions. We performed sensitivity analyses by repeating the analysis of **Figure 3** after excluding recording sites from groups of cortical functional regions (**Supplementary Figure 6**; groups of regions were: Auditory, Limbic, Visual + Ventral, Ventromedial Prefrontal + Lateral Prefrontal + Executive, and Dorsal + Action). The significant decrease in *D*_E_ during states of reduced consciousness was observed in all cases, regardless of which regions were eliminated. These results indicate that the changes in network structure associated with transitions into states of reduced consciousness could not be explained by connectivity changes of any single brain region. Thus, anesthesia and sleep are associated with global reorganization of cortical networks.

### Geometric correlates to changes in effective dimensionality

We’ve shown that *D*_E_ indexes entry and exit from states of reduced consciousness based on only the spectrum (eigenvalues) of **P_symm_**. The observed changes in *D*_E_ indicate a reorganization of brain networks; however, because there is no unique mapping between a spectrum and a network, changes in spectrum do not identify the specific features of this reorganization. To gain insight into these features, we can apply the next steps in DME analysis and consider data in the embedding space defined by the spectral decomposition of **P_symm_**.

A simple toy model is useful in this regard (**Figure 4**). We simulated a modular network consisting of five regions, with nine nodes in each region. Two types of connectivity were present in the model: 1) uniform random connectivity linking nodes regardless of region, and 2) stronger within-region connectivity imposed on this nonspecific random connectivity. The strength of within-region connectivity was varied from weak (**Figure 4a**, left column) to strong (**Figure 4a**, right column), corresponding to an increasingly modular organization of the network. This increase in modular organization was associated with more peaked eigenvalue spectra and decreased *D*_E_ (insets in **Figure 4a**). DME conveys the functional geometry of these changes in community structure by mapping the data into a lower dimensional embedding space using the eigenfunctions and eigenvalues of **P_symm_** (**Figure 4b**). Nodes that are connected similarly to the rest of the network are mapped to nearby locations in the embedding space, indicating their functional similarity. A more modular network organization results in more tightly clustered nodes within each region; the neural responses of this more modular network would exhibit reduced differentiation. This is easily illustrated by considering the extreme case (right), in which the nodes within each region are so tightly coupled as to render them nearly equivalent, essentially transforming the original 40-node network into a 5-node network with a vastly reduced repertoire of possible network states. In addition, regions become more distinct and more distant from each other as modularity increases, corresponding to a decrease in functional integration across the whole network.

**Figure 4.**
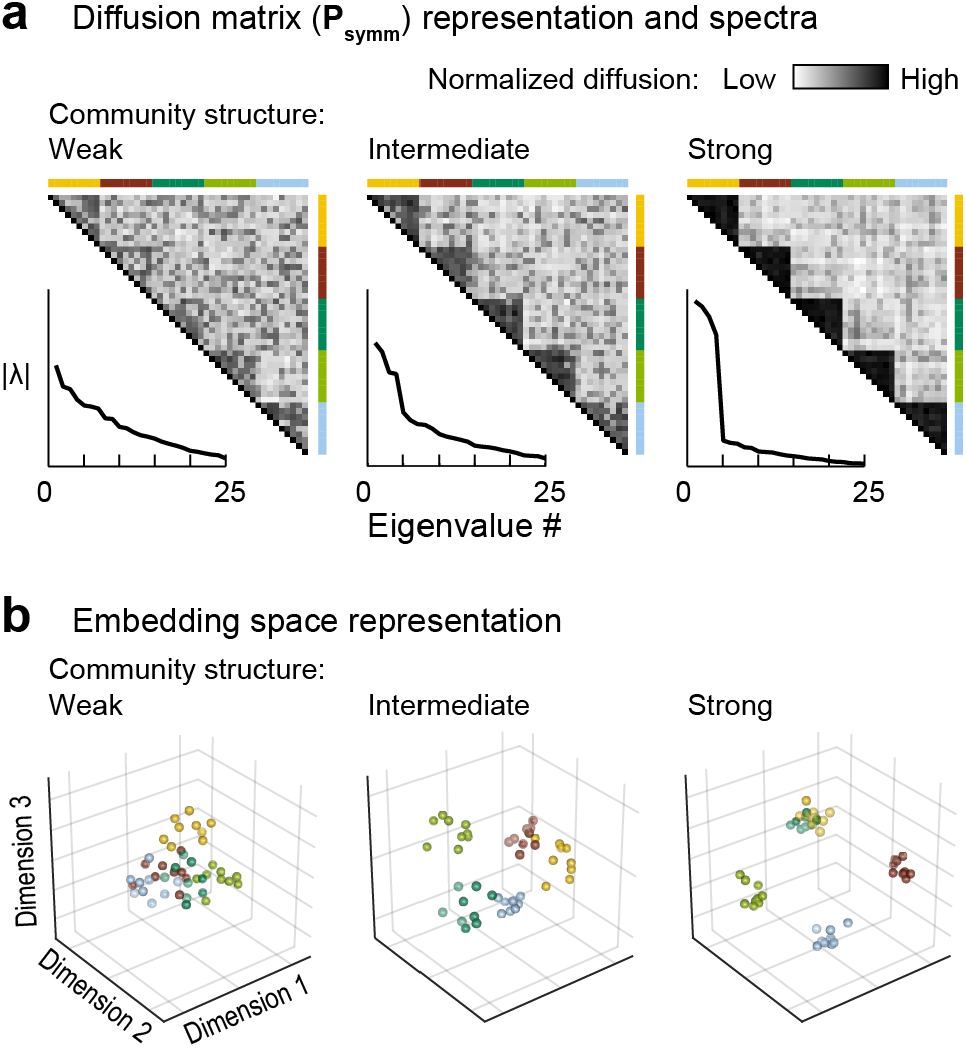
Toy example showing effects of stronger community structure on embeddings. **A**: Diffusion matrix (**P_symm_**) representation of weak, intermediate, and strong community structure (left, middle and right panel, respectively). Insets depict **P_symm_** spectra. *D*_E_ = 0.44, 0.35, 0.18, respectively. **B**: Embedding space representation of weak, intermediate, and strong community structure (left, middle and right panel, respectively). Mean centroid distance = 0.35, 0.42, 0.50, respectively.

We observed similar changes in embeddings of functional connectivity data derived from intracranial recordings (**Figure 5**). Data recorded during states of reduced consciousness during propofol anesthesia (e.g., U; **Figure 5a**) or sleep (e.g., N2 or N3; **Figure 5b**) were more ‘clumpy’ in embedding space and regional clusters of nodes moved farther apart from each other, suggesting an increase in modular organization.

**Figure 5.**
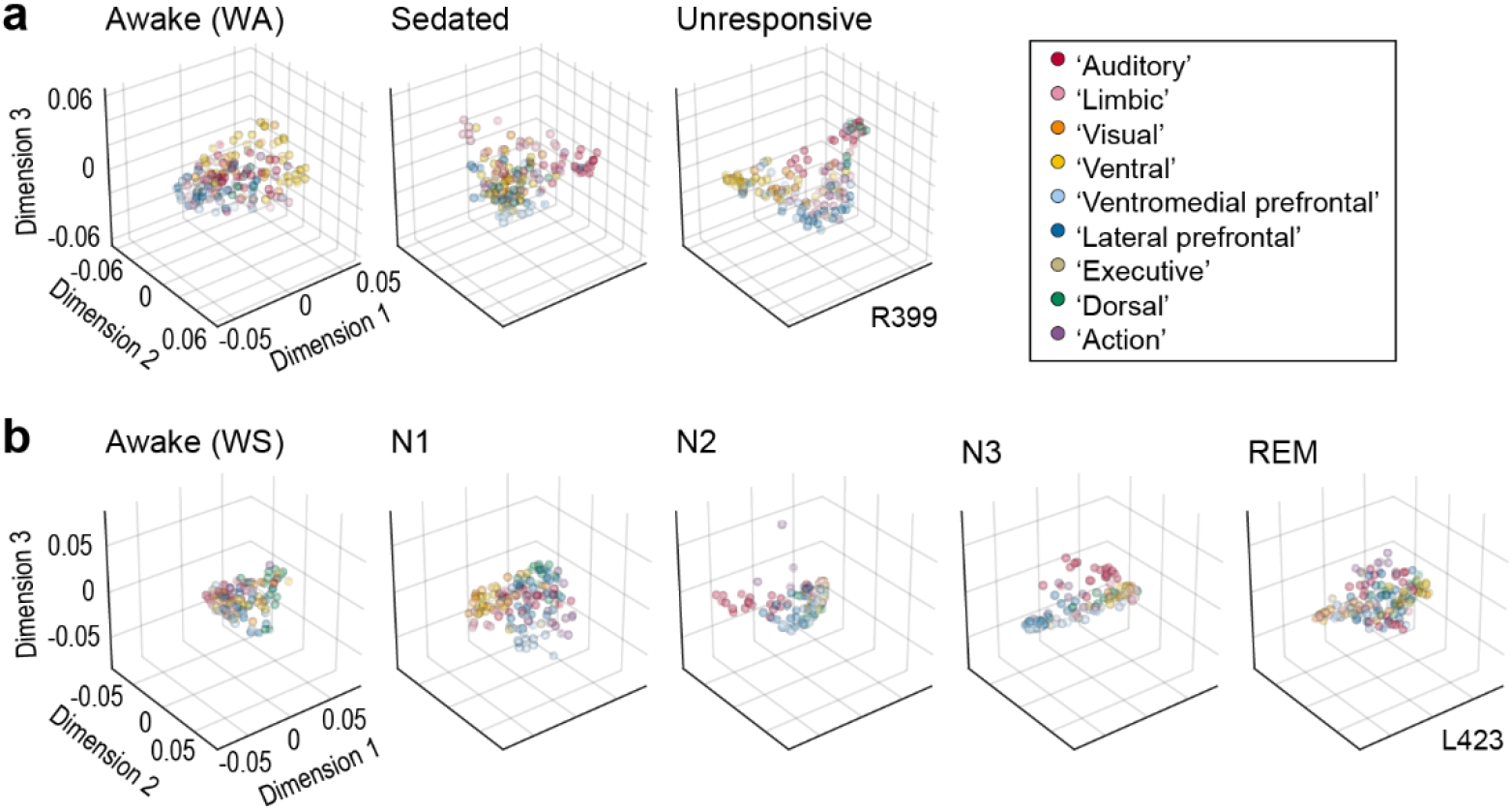
Changes in functional geometry during anesthesia and sleep in example participants. **A:** Arrangement of recorded data in embedding space (first three dimensions) during anesthesia in an example participant (R399). Each symbol represents an individual recording site. Colors indicate assignment to functional regions (legend). **B:** Similar to **a**, but for a second participant (L423) during sleep.

We quantified these effects both with and without *a priori* assignments of electrodes to labeled clusters. Using labels from the nine regions illustrated in **Supplementary Figure 1c**, we assessed changes in cluster organization within embeddings. We measured inter-cluster distances between cluster centroids and regional grouping of nodes using an index of cluster quality (Calinski-Harabasz index) calculated as the ratio of between-cluster and within-cluster dispersion. We also considered the position of nodes in embedding space relative to their neighboring nodes without *a priori* assignments to functional regions. The analysis is illustrated in **Supplementary Figure 7**. For each node, we calculated a normalized ‘local distance’ as the average pairwise distance to the 5^th^-percentile closest nodes normalized to the median distance to all nodes. Much like cluster quality this measure captures combined aspects of differentiation (distance among similar nodes) and integration (distance between dissimilar nodes), but without requiring label assignments.

Systematic changes in all three measures were observed across stages of anesthesia (**Figure 6a**, example participant). In U, inter-cluster distances increased, cluster quality improved, and local distances decreased. These results were consistent across subjects (likelihood ratio test for omitting state: inter-cluster distance χ^2^(2) = 37.8, p < 0.0001; cluster quality χ^2^(2) = 22.8, p < 0.0001; local distance χ^2^(2) = 47.9, p < 0.0001; see pairwise comparisons in **Supplementary Table 3**). Accordingly, effective dimensionality was negatively correlated with inter-cluster distance and cluster quality, and positively correlated with local distance (**Figure 6b**). Similar relationships with sleep stage were observed in an example participant (**Figure 6c**) and across participants (likelihood ratio test for omitting state: inter-cluster distance χ^2^(4) = 42.7, p < 0.0001; cluster quality χ^2^(4) = 22.5, p = 0.00016; local distance χ^2^(4) = 81.6, p < 0.0001; see pairwise comparisons in **Supplementary Table 3**) and were correlated with effective dimensionality (**Figure 6d**).

**Figure 6.**
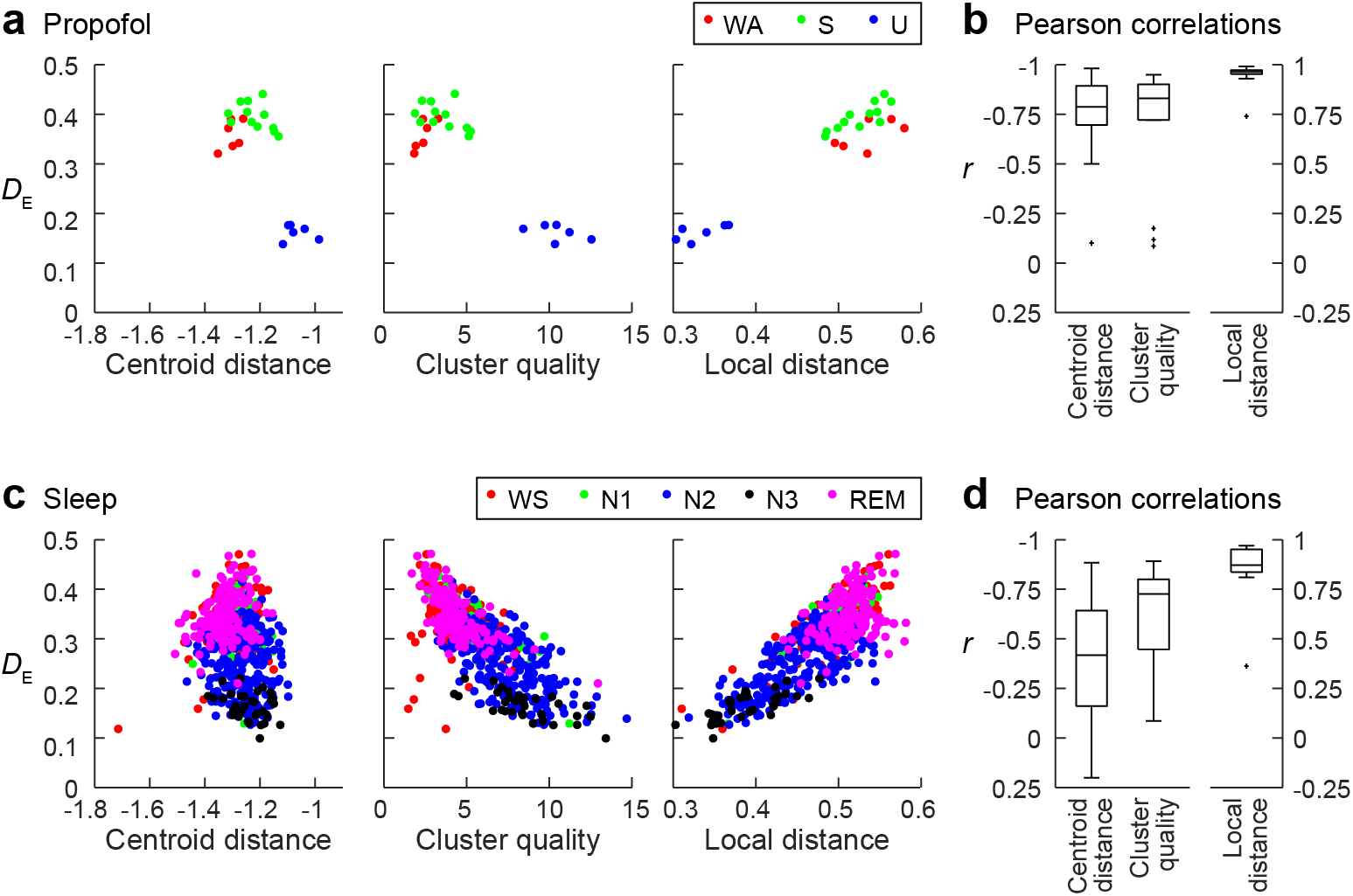
Changes in embedding geometry are correlated with effective dimensionality. During induction of anesthesia, inter-cluster distances, cluster quality, and local distance are state-dependent and therefore tend to correlate with effective dimensionality (**a-b**). **a:** Examples for a single participant (R399) where each point represents a 60-second segment of data. **b:** Pearson correlation coefficients across participants between *D*_E_ and each embedding measure. Centroid distance and cluster quality are negatively correlated with *D*_E_ and presented on a reversed axis. The strongest association with *D*_E_ is with local distance. Similar associations are observed during sleep (**c-d**). **c:** Single participant example (L423). **d:** Pearson correlation coefficients between embedding measures and *D*_E_ across participants.

For both sleep and anesthesia, the strongest correlations were observed between local distance and effective dimensionality. Local distance captures the reorganization in embedding space, and effective dimensionality allows for tracking changes in anesthesia or sleep stage, both without relying on *a priori* assumptions about the data.

## Discussion

### Summary of findings

Identifying changes in the brain that underlie LOC and ROC is a fundamental and unresolved question in neuroscience. Previous work indicates that consciousness is a property specific to the organizational structure of brain networks^1,46^. In this context, LOC and ROC should involve changes in this organization that are not tied to specific precipitating factors, such as the concentration of an exogenous anesthetic agent or the activity patterns in specific subcortical sleep and arousal centers. Building on our previous work^7^, we distinguished responsiveness under anesthesia (S) from unresponsiveness (U) to control for drug effects not specific to LOC. Although S is likely a stage of fluctuating arousal, we expected that participants were conscious a greater percentage of time compared to the unresponsive state U. Similarly, we expect that participants spent the majority of time conscious but drowsy or having conscious experiences in the form of dreaming during N1 and REM; this contrasts with N2 and N3, which are characterized by lower probability of dreaming and higher probability of unconsciousness^8^. We present a novel analytical framework that links changes in the organization of cortical networks with changes in arousal and awareness during anesthesia and sleep. We quantified these changes using *D*_E_, the effective dimensionality of a matrix derived from the functional connectivity matrix. *D*_E_ is robust to the choice of connectivity threshold used to construct the network graph, and is computationally efficient to calculate based on short data segments. *D*_E_ is also attractive because it is easily understood geometrically through its link to DME analysis.

### Changes in functional geometry reflect changes in complexity, differentiation, and integration

*D*_E_ is related to spatial complexity, in that fewer dimensions are required to represent a less complicated network. Thus, the results presented here are consistent with the decreased spatial complexity and smaller repertoire of distinct network configurations reported during LOC^47–51^. As shown in **Figure 4**, changes in simulated network modularity are reflected by changes in *D*_E_: when nodes are more tightly connected within each sub-network relative to connections to other sub-networks, *D*_E_ decreases (We note that there is not a strict relationship between *D*_E_ and modularity, as it is theoretically possible to create a biologically unrealistic network with no modularity but low *D*_E_.) In embedding space, this increase in modularity is reflected in increased cluster quality and in the shift to more local connectivity (**Figure 6**). Thus, the observation that *D*_E_ decreases during both anesthesia and sleep (**Figure 3**) links the results presented here with previous results derived from fMRI showing increased modularity during NREM sleep^27^. Reported decreases in within-network connectivity during anesthesia^32,33^ are harder to reconcile with increased modularity, though this could reflect differences in the spatial scale of the analyses.

Proximity in embedding space corresponds to similarity in functional connectivity to the rest of the network; during states of reduced consciousness, recording sites become closer to their nearest neighbors (less distinguishable) in embedding space, suggesting reduced differentiation of their activity patterns. Consistent with these results, perturbational complexity, a measure of differentiation and a well-validated measure of level of consciousness^24^, shows consistent decreases during anesthesia and sleep^34–37^. Similar results have been obtained using spatiotemporal complexity derived from resting state activity^38,39,52^.

The results presented here also speak to functional integration across the network, previously reported to decrease during anesthesia^40,53^ (though interestingly not during NREM sleep^27^). This is mostly easily visualized in embedding space, where functional regions tend to move farther apart during states of reduced consciousness (**Figure 6**). Decreased functional integration and differentiation during sleep and anesthesia likely play a role in reducing network efficiency during anesthesia and in disorders of consciousness^23,54,55^.

### Dynamics of network transitions

The dynamics of network transitions are becoming a rich vein of inquiry for understanding the neural basis of consciousness ^33,56^. Although these dynamics were not a focus of the current study, the framework presented here readily lends itself to their exploration. For example, simple clustering can distinguish integrated from segregated network states in resting state functional connectivity derived from fMRI data and enable exploration of the dynamics of state transitions during resting state and cognitive tasks^57,58^. These dynamics are altered under anesthesia, with a shift towards greater time spent in the segregated state and decreases in network complexity and information capacity^53,59^.

Although we have divided stages of anesthesia and sleep into two categories, one of reduced consciousness and the other of relatively intact consciousness, this is clearly an oversimplification. These stages of anesthesia (S, U) and sleep are undoubtedly superpositions of the more generally relevant states of unconsciousness, disconnected consciousness (i.e., dreaming), and connected consciousness. Stages of reduced consciousness (U, N2, N3) are likely dominated by segments of unconsciousness, but also include periods of disconnected consciousness^8,9^. Similarly, S and N1 are likely mixtures of connected consciousness, disconnected consciousness, and unconsciousness. This continuum is reflected in the smoothly varying changes across stage in *D*_E_ and other metrics presented here.

### Theories of consciousness

Central to the ongoing debate about the neural correlates of consciousness are their loci in the brain^14,15^. Global Neuronal Workspace Theory^46^ places prefrontal cortex and its connections with parietal regions central to these correlates, whereas Integrated Information Theory (IIT;^1^ sites these correlates in the ‘back’ of the brain, a region spanning temporal, occipital, and parietal cortex. Although clinical considerations precluded an exhaustive and invariant sampling of brain regions in our cohort of participants, our results indicate that transitions into and out of states of reduced consciousness involve a global network reorganization rather than relying on specific regions (**Supplementary Figure 6**). However, observation of global network changes during anesthesia and sleep does not exclude the possibility that local changes in key regions (i.e., prefrontal or parietal cortex) are sufficient to cause loss of consciousness. Additionally, even global cortical changes are likely coordinated by small brain areas with broad reach, such as central lateral thalamus^60^.

Previous work investigating mechanisms of anesthesia focused on disruptions in connectivity, especially feedback cortico-cortical connectivity. Studies in both human subjects and animal models showed reduced feedback connectivity at doses of anesthetics causing loss of consciousness^18,19,28,61^. These data are consistent with the Global Neuronal Workspace Theory, in which feedback from prefrontal cortex to wide areas of the brain is critical for conscious experience. They also support a model based on predictive processing in which consciousness relies on active comparisons between internally-generated expectations and observed sensory information^62,63^. However, several findings are not easily reconciled with these models, including reports of increases in connectivity^25–27^ and the reported suppression of feedforward connectivity^29^.

By focusing on network reorganization during states of reduced consciousness, we shift the focus beyond pathway-specific changes during LOC and ROC, and explore how local and global changes in connectivity combine to disrupt both differentiation and integration in the unconscious brain. However, we note that functional integration is not the same as information integration central to IIT as the latter distinguishes causal from non-causal interactions. Indeed, it is not possible to ascertain information integration using purely observational data. However, theoretical work has shown that differentiation can be used to establish an upper bound on integrated information^64^. Thus, the results presented here, when viewed through the lens of differentiation, are consistent with a decrease in information integration during reduced states of consciousness.

### Caveats & limitations

We focused here on a specific functional connectivity measure (orthogonalized power envelope correlations) and a specific frequency band (gamma). We focused on this band and this measure because our previous work demonstrated its utility for performing DME analysis^43^. However, we also note that the results of that paper were robust to choice of frequency band, and expect that the results presented here would also not depend strongly on that choice. Gamma band connectivity is also strongly related to connectivity derived from fMRI^65^, allowing comparisons to the body of work relying on neuroimaging for exploring changes in network organization during anesthesia and sleep.

Because participants in the current study had a neurological disorder, they may not be entirely representative of a healthy population. This caveat is inherent to all human intracranial electrophysiology studies, as discussed previously (e.g., Banks et al., 2020, 2022). However, we note that results presented in this study were consistent across participants with different seizure foci, clinical histories, and drug regimens. In addition, it is possible that seizures, AED use, and the hospital environment may affect sleep and sensitivity to anesthesia. Broadly, seizures can disrupt sleep architecture^66–68^. However, participants were monitored for seizure activity during the sleep recording session, and in the one participant with overnight seizures (L403), data collected after seizures began were excluded. Similarly, although AEDs are reported to alter the structure of sleep^69^, participants had discontinued their AEDs before collection of overnight sleep data, reducing the effect of AEDs on sleep data in this cohort. The quality and structure of sleep may also have been affected by the hospital environment, possibly contributing to the absence of N3 sleep in three participants. Because we had sufficient representation of all studied sleep stages in the cohort (see **Supplementary Table 1**), the effect of AEDs or the environment on the likelihood of entering a particular stage was not a confound. Similarly, while the use of AEDs could lead to a reduction of the dose of propofol required to achieve surgical level of general anesthesia^70^, the present study relied on behavioral assessment of arousal. Thus, the definition of stages of anesthesia was not affected by factors secondary to the participants’ history of epilepsy.

### Future directions

The iEEG results presented here support a model in which altered differentiation and functional integration of cortical networks underlie changes in consciousness, and suggest that the analytical framework presented here could contribute to understanding the neural correlates of consciousness. Next steps should include recapitulation of these results using non-invasive methods during anesthesia and sleep, and in patients with disorders of consciousness. Extending this analysis to scalp electroencephalography in particular would enhance the translational relevance of these findings. Assessments in clinical settings often require monitoring of consciousness in real time. Accordingly, tracking of the dynamics of *D*_E_ and of data in embedding space will enable identification of rapid changes in brain state underlying transitions between drowsiness, disconnected consciousness, and unconsciousness. Finally, extending DME analysis to apply to effective connectivity would enable more thorough investigation of causal structure theories of consciousness such as IIT.

## Materials and Methods

### Participants

The study was carried out in 21 neurosurgical patients (8 female; age 18-54 years old, median age 34 years old) diagnosed with medically refractory epilepsy. The patients were undergoing chronic invasive electrophysiological monitoring to identify seizure foci prior to resection surgery (**Supplementary Table 1**). Research protocols aligned with best practices recently aggregated in^71^ and were approved by the University of Iowa Institutional Review Board and the National Institutes of Health; written informed consent was obtained from all participants. Research participation did not interfere with acquisition of clinically necessary data, and participants could rescind consent for research without interrupting their clinical management. All participants underwent neuropsychological assessment prior to electrode implantation, and none had cognitive deficits that would impact the results of this study. The participants were tapered off their antiepileptic drugs during chronic monitoring when resting state data were collected.

### Experimental procedures

#### Pre-implantation neuroimaging

All participants underwent whole-brain high-resolution T1-weighted structural MRI scans before electrode implantation. The scanner was a 3T GE Discovery MR750W with a 32-channel head coil. The T1 scan (3D FSPGR BRAVO sequence) was obtained with the following parameters: FOV = 25.6 cm, flip angle = 12 deg., TR = 8.50 ms, TE = 3.29 ms, inversion time = 450 ms, voxel size = 1.0 × 1.0 × 0.8 mm.

#### iEEG recordings

iEEG recordings were obtained using either subdural and depth electrodes, or depth electrodes alone, based on clinical indications. Electrode arrays were manufactured by Ad-Tech Medical (Racine, WI). Subdural arrays, implanted in 14 participants out of 21, consisted of platinum-iridium discs (2.3 mm diameter, 5-10 mm inter-electrode distance), embedded in a silicon membrane. Stereotactically implanted depth arrays included between 4 and 12 cylindrical contacts along the electrode shaft, with 5-10 mm inter-electrode distance. A subgaleal electrode, placed over the cranial vertex near midline, was used as a reference in all participants. All electrodes were placed solely on the basis of clinical requirements, as determined by the team of epileptologists and neurosurgeons^72^.

Resting-state (RS) data were recorded during overnight sleep (*N* = 15 participants) and during induction of propofol anesthesia (*N* = 14 participants). In 8 participants, both sets of data were recorded, with sleep data collected first, followed several days later by anesthesia data.

#### Sleep recordings

Resting state iEEG, EEG, and video data were collected in the dedicated, electrically shielded suite in The University of Iowa Clinical Research Unit while the participants lay in the hospital bed. Sleep data were collected 7.5 +/- 1.1 days [range 6 – 9] after iEEG electrode implantation surgery. Data were recorded using a Neuralynx Atlas System (Neuralynx Inc., Bozeman, MT), amplified, filtered (0.1–500 Hz bandpass, 5 dB/octave rolloff), and digitized at a sampling rate of 2000 Hz.

Stages of sleep were defined manually using facial EMG and scalp EEG data based on standard clinical criteria (Berry et al., 2017) independently by two individuals who participate in the inter-scorer reliability program of the American Academy of Sleep Medicine: a licensed polysomnography technologist, certified by the Board of Registered Polysomnography Technologists, and a physician certified in Sleep Medicine by the Accreditation Council for Graduate Medical Education. The final staging report was agreed upon by the two scorers after a collaborative review. Scalp and facial electrodes were placed by an accredited technician, and data were recorded by a clinical acquisition system (Nihon Kohden EEG-2100) in parallel with research data acquisition. Facial electrodes were placed following guidelines of the American Academy of Sleep Medicine (Berry et al., 2017) at the left and right mentalis for EMG, and adjacent to left and right outer canthi for EOG. EEG was obtained from electrodes placed following the international 10-20 system at A1, A2, F3, F4, O1, and O2 in all participants, with the following additional electrodes: C3 and C4 in all participants but R376; E1 and E2 in L372 and R376; Cz and Fz in L409, L423, and L585; F7 in L585; F8 in L423 and L585. All participants had periods of N1 and N2 sleep identified; 12 out of 15 had N3 sleep periods and 12 out of 15 had REM. One participant (L403) experienced multiple seizures in the second half of the night; those data were excluded from analysis. The durations of recordings for each sleep stage in each participant are provided in **Supplementary Table 1**.

#### Anesthesia data

Resting state data were collected in the operating room during induction of propofol anesthesia just prior to electrode removal and seizure focus resection surgery. Data acquisition was controlled by a TDT RZ2 real-time processor (Tucker-Davis Technologies, Alachua, FL) in participants R369 through L460 and by a Neuralynx Atlas System in participants L514 and L585. Recorded data were amplified, filtered (0.7–800 Hz bandpass, 12 dB/octave rolloff for TDT-recorded data; 0.1–500 Hz bandpass, 5 dB/octave rolloff for Neuralynx-recorded data), and digitized at a sampling rate of 2034.5 Hz (TDT) or 2000 Hz (Neuralynx). Although no specific instructions were given about keeping eyes open or closed, participants were observed to have eyes closed during nearly all resting state recordings. Data were recorded in 3-4 blocks (duration 3-6 minutes each), interleaved with auditory stimulus paradigms related to other studies (e.g.,^73,74^). Data were collected during an awake baseline period and during infusion of increasing doses of propofol (50 – 150 μg/kg/min; **Supplementary Figure 2**).

Awareness was assessed using the Observer’s Assessment of Alertness/Sedation (OAA/S) scale (Chernik et al., 1990). Bispectral index (BIS) (Gan et al., 1997) was measured using BIS Complete 4-Channel Monitor (Medtronic plc, Minneapolis, MN), but was not used in the analyses presented in this study. OAA/S was assessed just before and just after collection of each resting state data block. Data segments were assigned labels corresponding to one of three stages of the anesthesia experiment: wake (WA; i.e., pre-drug) and two levels of anesthesia: sedated but responsive to command (S; OAA/S ≥ 3) and unresponsive (U; OAA/S ≤ 2) (Nourski et al., 2018a) (**Supplementary Figure 2**).

In 6 of 14 participants, OAA/S values crossed the boundary between S and U over the course of the resting state block (e.g. resting state block #1 in participant L372; see **Supplementary Figure 2**). In these cases, only the first and last 60-second segments of the block were analyzed; data from the first segment were labeled S, and data from the second segment were labeled U. Data in the intervening segment were not assigned an anesthesia stage label and were not used in the analysis. The durations of recordings used in the analyses for each stage and each participant during the anesthesia experiment are provided in **Supplementary Table 1**.

### Data analysis

#### Anatomical reconstruction and ROI parcellation

Localization of recording sites and their assignment to ROIs relied on post-implantation T1-weighted anatomical MRI and post-implantation computed tomography (CT). All images were initially aligned with pre-operative T1 scans using linear coregistration implemented in FSL (FLIRT)^75^. Electrodes were identified in the post-implantation MRI as magnetic susceptibility artifacts and in the CT as metallic hyperdensities. Electrode locations were further refined within the space of the pre-operative MRI using three-dimensional non-linear thin-plate spline warping^76^, which corrected for post-operative brain shift and distortion. The warping was constrained with 50-100 control points, manually selected throughout the brain, which were visually aligned to landmarks in the pre- and post-implantation MRI.

To sort recording sites for presentation of diffusion matrices and for assessment of centroid distances and clustering, recording sites were assigned to one of 58 ROIs organized into 9 functional regions (see **Figure 1**, **Supplementary Figure 1**, **Supplementary Table 2**)^43^ based on anatomical reconstructions of electrode locations in each participant. For subdural arrays, ROI assignment was informed by automated parcellation of cortical gyri^77,78^ as implemented in the FreeSurfer software package. For depth arrays, it was informed by MRI sections along sagittal, coronal, and axial planes. Subcortical recording sites, recording sites identified as seizure foci or characterized by excessive noise, and depth electrode contacts localized to the white matter or outside brain, were excluded from analyses and are not listed in **Supplementary Table 2**.

#### Preprocessing of iEEG data

Analysis of iEEG data was performed using custom software written in MATLAB Version 2021b programming environment (MathWorks, Natick, MA, USA). After initial rejection of recording sites identified as seizure foci, several automated steps were taken to exclude recording channels and time intervals contaminated by noise. First, channels were excluded if average power in any frequency band (broadband, delta, theta, alpha, beta, gamma, or high gamma; see below) exceeded 3.5 standard deviations of the average power across all channels for that participant. Next, transient artifacts were detected by identifying voltage deflections exceeding 10 standard deviations on a given channel. A time window was identified extending before and after the detected artifact until the voltage returned to the zero-mean baseline plus an additional 100 ms buffer before and after. High-frequency artifacts were also removed by masking segments of data with high gamma power exceeding 5 standard deviations of the mean across all segments. Only time bins free of these artifact masks were considered in subsequent analyses. Artifact rejection was applied across all channels simultaneously so that all connectivity measures were derived from the same time windows. Occasionally, particular channels survived the initial average power criteria yet had frequent artifacts that led to loss of data across all the other channels. There is a tradeoff in rejecting artifacts (losing time across all channels) and rejecting channels (losing all data for that channel). If artifacts occur on many channels, there is little benefit to excluding any one channel. However, if frequent artifacts occur on one or simultaneously on up to a few channels, omitting these can save more data from other channels than those channels contribute at all other times. We chose to optimize the total data retained, channels × time windows, and omitted some channels when necessary. To remove shared signals unlikely to derive from brain activity, data from retained channels were high-pass filtered above 200 Hz, and a spatial filter was derived from the singular value decomposition omitting the first singular vector. This spatial filter was then applied to the broadband signal to remove this common signal.

For connectivity analysis, the orthogonalized gamma band (30-70 Hz) power envelope correlation^45^ was used. This measure avoids artifacts due to volume conduction by discounting connectivity near zero phase lag. Data were divided into 60-second segments, pairwise connectivity estimated in each segment, and then connectivity estimates averaged across all segments for that participant.

Envelope correlations were estimated for each data segment and every recording site as in^45^, except time-frequency decomposition was performed using the demodulated band transform^79^, rather than wavelets. Gamma power at each time bin was calculated as the average (across frequencies) log of the squared amplitude. For each pair of signals *X* and *Y*, one was orthogonalized to the other by taking the magnitude of the imaginary component of the product of one signal with the normalized complex conjugate of the other:

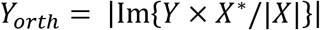

Both signals were band-pass filtered (0.2 – 1 Hz), and the Pearson correlation calculated between signals. The process was repeated by orthogonalizing in the other direction and the overall envelope correlation for a pair of recording sites was the average of the two Pearson correlations.

Prior to diffusion map embedding, connectivity matrices were thresholded by saving at least the top third (rounded up) connections for every row, as well as their corresponding columns (to preserve symmetry). We also included any connections making up the minimum spanning tree of the graph represented by the elementwise reciprocal of the connectivity matrix to ensure the graph is connected.

To confirm that the results presented here did not depend on the specific threshold chosen, two additional thresholds were tested: 1) a more strict procedure, using the same procedure as above except saving only the top 10%, or 2) a more permissive procedure, only thresholding out negative correlations.

#### Diffusion map embedding

See Banks et al. (2022)^43^ for details about DME. In brief, cosine similarity was applied to the functional connectivity matrix (here orthogonalized power envelope correlations) to yield the similarity matrix **K =** [*k*(*i*,*j*)], which was normalized by degree to yield a diffusion matrix **P = D**^−1^**K**, where **D** is the degree matrix, i.e. the diagonal elements of 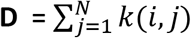, where *N* is the number of recording sites, and the off-diagonal elements of **D** are zero. If the recording sites are conceptualized as nodes on a graph with edges defined by **K**, then **P** can be understood as the transition probability matrix for a ‘random walk’ or a ‘diffusion’ on the graph (see^42,80^). DME consists of mapping the recording sites into an embedding space using an eigendecomposition of **P**,

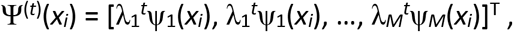

where ψ_*j*_ are the eigenvectors of **P**. The parameter *t* is the number of time steps in that random walk; here, we fix *t* = 1. DME can be implemented alternatively based on a symmetric version of diffusion matrix **P_symm_** = **D**^−0.5^**KD**^−0.5^. Basing DME on **P_symm_** has the advantage that the eigenvectors of **P_symm_** form an orthogonal basis set (unlike the eigenvectors of **P**), providing some additional convenience mathematically that is beyond the scope of this paper^42^. Additionally, the eigenvalues of **P** and **P_symm_** are identical.

#### Effective dimensionality

We used effective dimensionality (*D_E_*)^81^, a graph theoretic measure of network complexity, to characterize the shape of the spectrum of **P_symm_**, or equivalently the complexity of its community structure. *D_E_* was calculated from the eigenvalue spectrum | λ_i_| of **P_symm_** and normalized to the total number of dimensions (*N*; equal to the number of recording sites) as 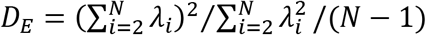. The first dimension for which λ_1_ = 1 is skipped. *D_E_* gives information about how data is distributed in *N* dimensions (where *N* is the number of recording sites). *D_E_* = 1 for a random graph, as the data are distributed equally in every dimension and the spectrum is flat. A graph with structure, e.g., nodes that connect to each other more than the rest of the graph, has a peaked spectrum and *D_E_* <1.

#### Dimensionality reduction via low rank approximations to P_symm_

When calculating distances or evaluating clustering in embedding space, we used a low rank approximation, discarding dimensions associated with small eigenvalues that are likely dominated by noise. The choice of threshold for this procedure is somewhat arbitrary; we used an algorithm to identify the inflection point *k*_infl_ beyond which eigenvalues are small and decrease gradually^82^, and the number of dimensions retained set equal to *k*_infl_ – 1.

#### Clustering of functional regions in embedding space

ROIs were categorized into 9 functional regions based on analysis of resting state data from a different cohort of participants (Banks et al., 2022). A small number of sites in ROIs that were not used in the scheme in Banks et al., 2022 were assigned to ROI clusters based on anatomical and functional criteria.

Two measures were used to quantify the arrangement of nodes in embedding space according to these functional regions. First, the distance between regions in embedding space was measured by the pairwise (by region) Euclidean distance between centroids (mean position across nodes within each region). Second, the Calinski-Harabasz index of cluster quality (the ratio of between-cluster variance to within-cluster variance;^83^) was used to quantify the extent to which nodes segregated in embedding space according to these pre-identified functional regions.

#### Local distance

To quantify the tendency of nodes to be functionally distinct from other nodes (or, conversely, to aggregate in embedding space and be less differentiated) without needing to rely on assignments of nodes to pre-defined ROIs or regional groupings, we defined a measure called ‘local distance’ as the mean Euclidean distance in embedding space from a given node to each of the 5% closest other nodes, divided by the median distance to all pairs of nodes.

#### Statistical modeling

All measures (*D_E_*, centroid distance, Calinski-Harabasz index, local distance) were computed for individual data segments, then averaged within each participant across all segments of the same behavioral state (WA, S, U, WS, N1, N2, N3, R). Linear mixed effects models were fit to these measures with behavioral state as a fixed effect and participant as a random effect; fit models were compared to a reduced model omitting the fixed effect for state using a likelihood ratio test. Pairwise planned contrasts were tested between WA-S, WA-U, and S-U for propofol experiments, and WS-N1, WS-N2, WS-N3, WS-R, N1-N2, N2-N3, N2-R and N3-R for sleep experiments; p-values were adjusted using a multivariate *t* distribution that accounts for correlations among tested hypotheses. Statistical analyses were performed in R version 4.2.1 using the packages lme4^84^ and emmeans^85^.

## Acknowledgements

This work was supported by the National Institutes of Health (grant numbers R01-DC04290, R01-GM109086, S10OD025025, UL1-RR024979). We are grateful to Haiming Chen, Brian Dlouhy, M. Eric Dyken, Phillip Gander, Christopher Garcia, Timothy Griffiths, Matthew Howard, William Mayner, Christopher Petkov, and Ariane Rhone for help with data collection, analysis, interpretation, and helpful comments on the manuscript.

## Data and code availability

Full data is available via a request to the Authors pending establishment of a formal data sharing agreement. Data required to reproduce figures from the manuscript and statistical analyses are provided with the software. Software is available at: https://zenodo.org/record/7320253 or https://doi.org/10.5281/zenodo.7320253

## Author contributions

Conceptualization: K.V.N., M.I.B.

Methodology: B.M.K., M.I.B, K.V.N.

Software: B.M.K., D.I.C.

Validation: B.M.K.

Formal Analysis: B.M.K., D.I.C., C.K.K., M.I.B.

Investigation: R.N.M., H.K., K.V.N, M.I.B.

Data Curation: B.M.K., C.K.K., K.V.N.

Writing – original draft: B.M.K., K.V.N., M.I.B.

Writing – Review & Editing: B.M.K., D.I.C., C.K.K., R.N.M., H.K., K.V.N, M.I.B.

Visualization: B.M.K, K.V.N.

Supervision: K.V.N., M.I.B.

Project Administration: K.V.N., M.I.B.

Funding Acquisition: K.V.N., M.I.B.

