## Supplemental Tables and Figures for "Consciousness is indexed by analogous cortical reorganization during sleep and anesthesia"

### Supplementary Information

Bryan M. Krause[1], Declan I. Campbell[1], Christopher K. Kovach[2], Rashmi N. Mueller[2,3], Hiroto Kawasaki[2], Kirill V. Nourski[2,4], Matthew I. Banks[1,5*]

[1]*Department of Anesthesiology, University of Wisconsin, Madison, WI, USA*
[2]*Department of Neurosurgery, The University of Iowa, Iowa City, IA 52242, USA*
[3]*Department of Anesthesia, The University of Iowa, Iowa City, IA 52242, USA*
[4]*Iowa Neuroscience Institute, The University of Iowa, Iowa City, IA 52242, USA*
[5]*Department of Neuroscience, University of Wisconsin, Madison, WI, USA*

***Corresponding author:**

Matthew I. Banks, Ph.D.
Professor
Department of Anesthesiology
University of Wisconsin
1300 University Avenue, Room 4605
Madison, WI 53706


**Keywords:**

Functional connectivity, intracranial electrophysiology, anesthesia, sleep

**Acknowledgements:**

This work was supported by the National Institutes of Health (grant numbers R01-DC04290, R01-GM109086, S10OD025025, UL1-RR024979). We are grateful to Haiming Chen, Brian Dlouhy, M. Eric Dyken, Phillip Gander, Christopher Garcia, Timothy Griffiths, Matthew Howard, William Mayner, Christopher Petkov, and Ariane Rhone for help with data collection, analysis, interpretation, and helpful comments on the manuscript.

### Supplementary Table 1. Subject demographics and summary of anesthesia and sleep data.

| Subject | Age | Sex | Clinical background | | | Recording duration (min.) | | | | | | | | | |
|---|---|---|---|---|---|---|---|---|---|---|---|---|---|---|---|
| | | | | | | Propofol | | | | Sleep | | | | | |
| | | | MRI | PET | Seizure focus | W | S | U | Total | W | N1 | N2 | N3 | REM | Total |
| R369 | 30 | M | R basal ganglia deep venous anomaly | R mesial temporal hypometabolism | R medial temporal | - | - | - | - | 98 | 5 | 10 | 0 | 0 | 113 |
| L372 | 34 | M | Normal | L mesial temporal hypometabolism | L temporal pole | 6 | 1 | 7 | 14 | 126 | 16 | 178 | 0 | 47 | 367 |
| R376 | 48 | F | L and R frontal white matter small vessel ischemic disease | Normal | R medial temporal | 6 | 6 | 6 | 18 | 148 | 12 | 285 | 42 | 47 | 534 |
| R384 | 38 | M | R mesial temporal sclerosis | R anterior and mesial temporal hypometabolism | R medial temporal | 6 | 0 | 12 | 18 | - | - | - | - | - | - |
| R394 | 24 | M | R anterior and medial temporal post-surgical change | R temporal hypometabolism | R amygdala (medial temporal) | 6 | 6 | 6 | 18 | - | - | - | - | - | - |
| R399 | 22 | F | Normal | R lateral and superior temporal hypometabolism | R temporal (uncertain medial & neocortical) | 5 | 1 | 7 | 13 | - | - | - | - | - | - |
| L400 | 59 | F | L mesial temporal sclerosis, L dorsolateral prefrontal volume loss | L hemisphere diffuse hypometabolism except occipital lobe | L amygdala (medial temporal) | 6 | 7 | 1 | 14 | - | - | - | - | - | - |
| L403 | 56 | F | Normal | R mesial temporal slight hypometabolism | L medial temporal | 6 | 1 | 7 | 14 | 111 | 30 | 141 | 0 | 1 | 283 |
| L405 | 19 | M | Left frontal encephalomalacia | L frontal hypometabolism | L frontal (posterior lateral) | 6 | 1 | 7 | 14 | - | - | - | - | - | - |
| L409 | 31 | F | L amygdala enlargement, L medial temporal choroidal fissure cyst | L mesial temporal hypometabolism | L medial temporal, L temporal pole | 6 | 7 | 1 | 14 | 165 | 46 | 189 | 9 | 82 | 491 |
| R413 | 22 | M | R medial temporal subtle FLAIR signal change | R anterior and medial temporal hypometabolism | right medial temporal | 6 | 6 | 9 | 21 | - | - | - | - | - | - |
| R418 | 25 | F | R posterior lateral ventral cortex developmental malformation | R temporal hypometabolism | R temporal (medial & lateral posterior cortex) | 6 | 12 | 6 | 24 | 132 | 34 | 253 | 34 | 150 | 603 |
| L423 | 51 | M | Normal | Normal | L medial temporal | 6 | 7 | 7 | 20 | 307 | 32 | 199 | 20 | 42 | 600 |
| L439 | 37 | M | Right parieto-occipital encephalomalacia, bilateral frontal R basal ganglia gliosis | Not done | R medial frontal (posterior dorsal) | - | - | - | - | 199 | 32 | 75 | 65 | 0 | 371 |
| L457 | 18 | M | Normal | Normal | L medial temporal | - | - | - | - | 282 | 10 | 180 | 70 | 77 | 619 |
| L460 | 52 | M | Normal | L medial temporal hypometabolism | L hippocampus (medial temporal) | - | - | - | - | 70 | 43 | 243 | 41 | 30 | 427 |
| L514 | 46 | M | L insula atrophy | Normal | L insula (anterior) | 6 | 0 | 6 | 12 | 74 | 67 | 338 | 10 | 39 | 528 |
| R524 | 18 | M | Normal | R anterior and medial temporal hypometabolism | R medial temporal, L medial temporal | - | - | - | - | 63 | 7 | 315 | 28 | 65 | 478 |
| R532 | 42 | F | Normal | R medial temporal hypometabolism | R ventral frontal (posterior) | - | - | - | - | 341 | 21 | 186 | 5 | 83 | 636 |
| R567 | 33 | M | Normal | Not done | R insula | - | - | - | - | 315 | 105 | 251 | 8 | 0 | 679 |
| L585 | 39 | F | L anterior temporal gray-white matter differentiation blurring | L anterior temporal hypometabolism | L medial temporal | 10 | 0 | 6 | 16 | 223 | 34 | 191 | 41 | 162 | 651 |

### Supplementary Table 2. Electrode coverage.

| ROI group | ROI | Propofol | | Sleep | |
|---|---|---|---|---|---|
| | | $N_{subjects}$ | $n_{sites}$ | $N_{subjects}$ | $n_{sites}$ |
| **'Auditory'** | HGPM | 13 | 64 | 12 | 67 |
| | HGAL | 9 | 29 | 8 | 26 |
| | PT | 11 | 35 | 9 | 38 |
| | STGP | 12 | 107 | 12 | 103 |
| | STGM | 13 | 60 | 11 | 53 |
| | STSU | 8 | 26 | 8 | 41 |
| **'Limbic'** | InsP | 11 | 31 | 11 | 31 |
| | InsA | 9 | 18 | 10 | 20 |
| | TP | 13 | 91 | 10 | 76 |
| | PHG | 13 | 41 | 9 | 24 |
| | Amyg | 9 | 26 | 8 | 23 |
| | Hipp | 6 | 13 | 9 | 23 |
| **'Visual'** | LingG | 6 | 9 | 3 | 5 |
| | Cun | 2 | 3 | 2 | 2 |
| | FG | 11 | 44 | 11 | 41 |
| | IOG | 3 | 3 | 1 | 1 |
| | ITGP | 10 | 34 | 8 | 22 |
| | ITGM | 11 | 34 | 10 | 35 |
| **'Ventral'** | PP | 11 | 25 | 10 | 23 |
| | STSL | 9 | 27 | 11 | 36 |
| | STGA | 9 | 24 | 8 | 20 |
| | MTGP | 13 | 135 | 12 | 113 |
| | MTGM | 13 | 87 | 12 | 79 |
| | MTGA | 11 | 42 | 12 | 50 |
| | ITGA | 11 | 51 | 10 | 48 |
| **'Ventromedial prefrontal'** | ACC | 7 | 9 | 6 | 9 |
| | fOperc | 5 | 7 | 2 | 4 |
| | OG | 14 | 127 | 14 | 138 |
| | GR | 11 | 21 | 10 | 15 |
| | FMG | 1 | 2 | 2 | 5 |
| | SubcG | 0 | 0 | 2 | 4 |
| **'Lateral prefrontal'** | IFGop | 12 | 42 | 12 | 38 |
| | IFGtr | 11 | 48 | 12 | 58 |
| | IFGor | 6 | 12 | 8 | 18 |
| | MFG | 11 | 135 | 12 | 122 |
| | SFG | 6 | 51 | 7 | 59 |
| | TFG | 9 | 31 | 8 | 20 |
| **'Executive'** | CingM | 6 | 11 | 9 | 24 |
| | PMC | 12 | 42 | 12 | 70 |
| | PCCpreCun | 0 | 0 | 2 | 7 |
| **'Dorsal'** | SMG | 12 | 90 | 11 | 109 |
| | AGP | 7 | 21 | 6 | 30 |
| | AGA | 10 | 46 | 10 | 58 |
| | MOG | 5 | 15 | 5 | 19 |
| **'Action'** | dPreCG | 11 | 37 | 11 | 45 |
| | dPostCG | 5 | 22 | 9 | 29 |
| | vPreCG | 13 | 50 | 14 | 62 |
| | vPostCG | 9 | 44 | 12 | 50 |
| | ParaCL | 0 | 0 | 1 | 2 |
| | SPL | 3 | 10 | 4 | 11 |
| | pOperc | 3 | 8 | 3 | 11 |
| **Total** | | 14 | 1932 | 15 | 2017 |

### Supplementary Table 3. Post-hoc pairwise comparisons.

| Measure | Contrast | Estimate | SE | df | *t* | *p* |
|---|---|---|---|---|---|---|
| **Mean $D_E$, propofol** | S - WA | -0.048 | 0.0165 | 23.9 | -2.92 | 0.020 |
| | U - WA | -0.15 | 0.0151 | 23.4 | -9.74 | <0.0001 |
| | S - U | 0.099 | 0.0165 | 23.9 | 6.01 | <0.0001 |
| **Mean $D_E$, sleep** | N1 - WS | -0.018 | 0.0122 | 50.0 | -1.47 | 0.55 |
| | N2 - WS | -0.086 | 0.0122 | 50.0 | -7.04 | <0.0001 |
| | N3 - WS | -0.13 | 0.0131 | 50.2 | -10.3 | <0.0001 |
| | R - WS | 0.0047 | 0.0131 | 50.2 | 0.362 | 1.00 |
| | N1 - N2 | 0.068 | 0.0122 | 50.0 | 5.57 | <0.0001 |
| | N2 - N3 | 0.049 | 0.0131 | 50.2 | 3.78 | 0.0032 |
| | N2 - R | -0.090 | 0.0131 | 50.2 | -6.91 | <0.0001 |
| | N3 - R | -0.14 | 0.0139 | 50.2 | -10.1 | <0.0001 |
| **Centroid distance, propofol** | S - WA | 0.075 | 0.0224 | 23.1 | 3.34 | 0.0077 |
| | U - WA | 0.18 | 0.0204 | 23.0 | 8.96 | <0.0001 |
| | S - U | -0.11 | 0.0224 | 23.1 | -4.83 | 0.0002 |
| **Centroid distance, sleep** | N1 - WS | 0.016 | 0.0115 | 50.0 | 1.02 | 0.82 |
| | N2 - WS | 0.059 | 0.0115 | 50.0 | 3.81 | 0.0028 |
| | N3 - WS | 0.12 | 0.0167 | 50.0 | 6.92 | <0.0001 |
| | R - WS | 0.021 | 0.0167 | 50.0 | 1.24 | 0.70 |
| | N1 - N2 | -0.043 | 0.0155 | 50.0 | -2.79 | 0.047 |
| | N2 - N3 | -0.056 | 0.0167 | 50.0 | -3.38 | 0.0097 |
| | N2 - R | 0.038 | 0.0167 | 50.0 | 2.30 | 0.14 |
| | N3 - R | 0.095 | 0.0177 | 50.0 | 5.36 | <0.0001 |
| **Cluster quality, propofol** | S - WA | 3.9 | 1.98 | 23.4 | 1.97 | 0.14 |
| | U - WA | 10.7 | 1.83 | 22.4 | 5.84 | <0.0001 |
| | S - U | -6.8 | 1.98 | 23.4 | -3.41 | 0.0064 |
| **Cluster quality, sleep** | N1 - WS | 1.9 | 1.71 | 50.3 | 1.09 | 0.78 |
| | N2 - WS | 5.7 | 1.71 | 50.3 | 3.33 | 0.011 |
| | N3 - WS | 8.1 | 1.84 | 50.9 | 4.38 | 0.0005 |
| | R - WS | 1.7 | 1.84 | 50.9 | 0.941 | 0.86 |
| | N1 - N2 | -3.8 | 1.71 | 50.3 | -2.24 | 0.16 |
| | N2 - N3 | -2.4 | 1.84 | 50.8 | -1.28 | 0.67 |
| | N2 - R | 4.0 | 1.84 | 50.9 | 2.16 | 0.19 |
| | N3 - R | 6.3 | 1.95 | 51.1 | 3.25 | 0.014 |
| **Local distance, propofol** | S - WA | -0.064 | 0.0192 | 23.2 | -3.31 | 0.0084 |
| | U - WA | -0.19 | 0.0177 | 22.2 | -10.8 | <0.0001 |
| | S - U | 0.13 | 0.0192 | 23.2 | 6.68 | <0.0001 |
| **Local distance, sleep** | N1 - WS | -0.019 | 0.0114 | 49.9 | -1.64 | 0.44 |
| | N2 - WS | -0.078 | 0.0114 | 49.9 | -6.82 | <0.0001 |
| | N3 - WS | -0.13 | 0.0123 | 50.1 | -10.9 | <0.0001 |
| | R - WS | -0.0058 | 0.0123 | 50.1 | -0.473 | 0.99 |
| | N1 - N2 | 0.059 | 0.0114 | 49.9 | 5.18 | <0.0001 |
| | N2 - N3 | 0.056 | 0.0123 | 50.1 | 4.54 | 0.0003 |
| | N2 - R | -0.072 | 0.0123 | 50.1 | -5.87 | <0.0001 |
| | N3 - R | -0.13 | 0.0130 | 50.1 | -9.82 | <0.0001 |

SE = standard error. df = degrees of freedom, adjusted by Satterthwaite method.

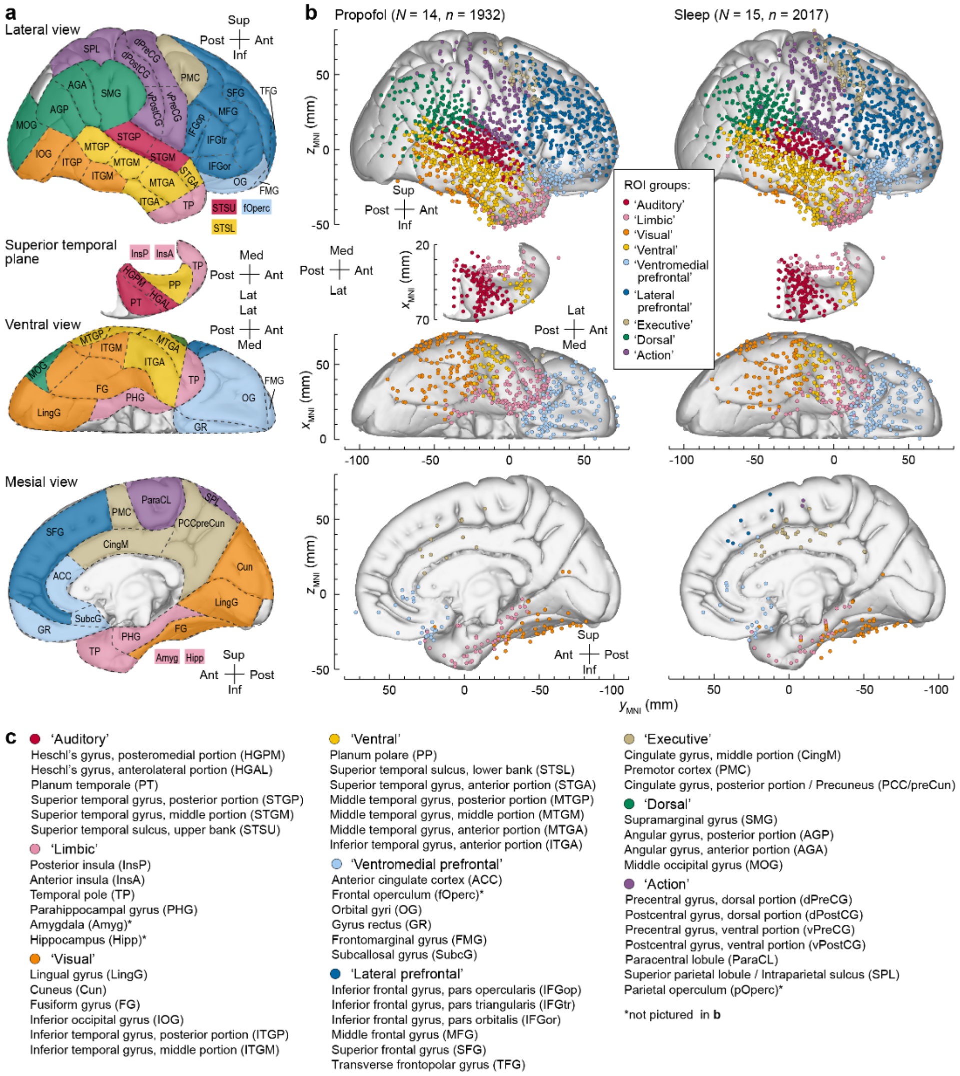


**Supplementary Figure 1. Regions of interest (ROI) and electrode coverage. a:** Region of interest (ROI) parcellation scheme used in the present study. **b:** Electrode coverage in all subjects that contributed to the propofol and sleep data sets (left and right column, respectively). Locations of recording sites, color-coded according to functional region, are plotted in MNI coordinate space and projected onto the right hemisphere of the MNI152 average template brain for spatial reference. Left hemisphere MNI *x*-axis coordinates ($x_{MNI}$) were multiplied by -1 to map them onto the right-hemisphere common space. Projection is shown on the lateral, top-down (STP), ventral and mesial views, aligned with respect to the $y_{MNI}$ coordinate (top to bottom rows). Recording sites over orbital, frontomarginal, inferior temporal gyrus and temporal pole are shown in both the lateral and the ventral view. Sites in fusiform, lingual, parahippocampal gyrus and gyrus rectus are shown in both the ventral and medial view. Sites in the amygdala, frontal operculum, hippocampus, and parietal operculum are not shown. **c:** ROI groups, ROIs, and abbreviations used in the present study.

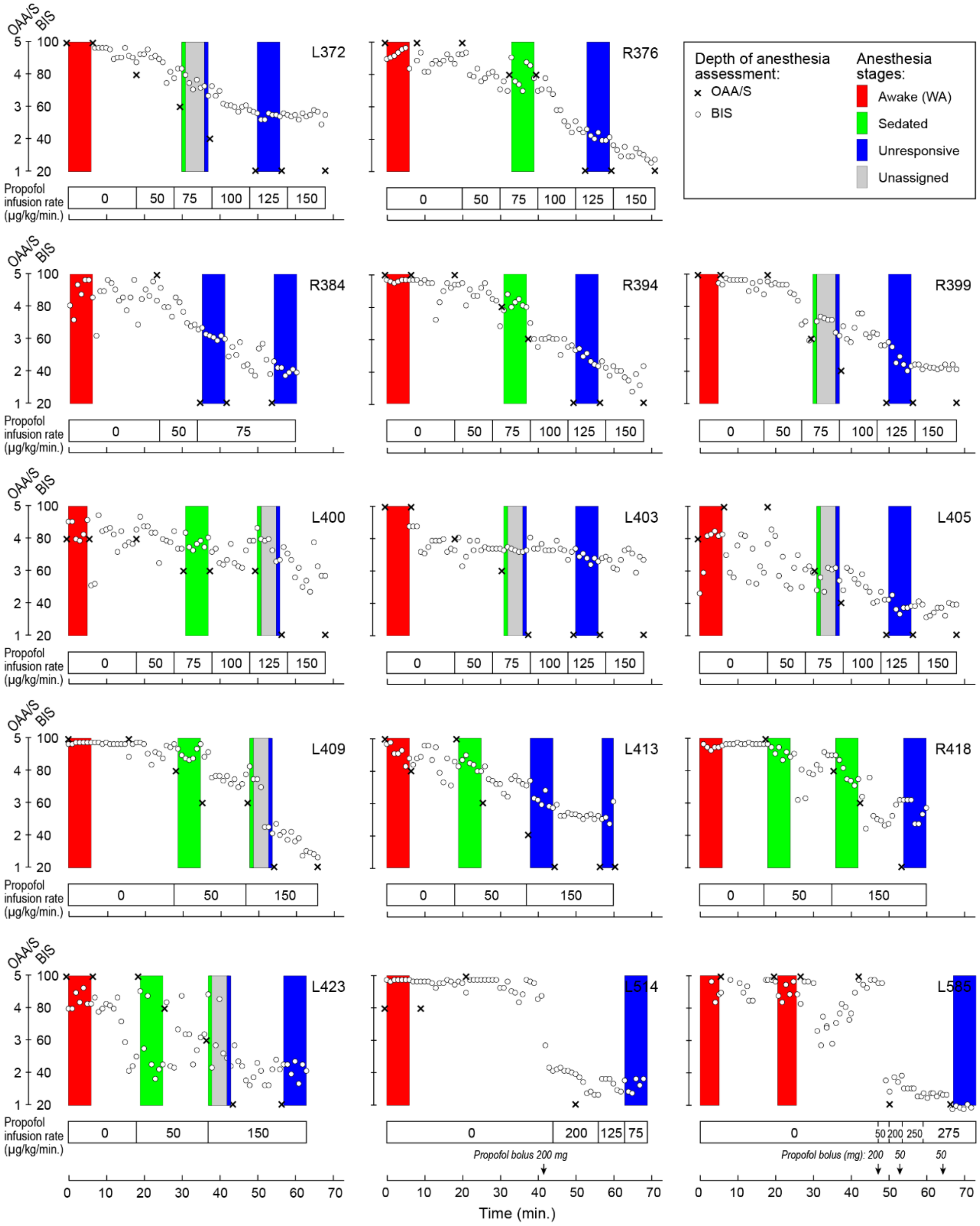


**Supplementary Figure 2. Induction of general anesthesia in all participants.** Observer's Assessment of Alertness/Sedation (OAA/S) scores (crosses) and bispectral index (BIS) values (open circles) are plotted as functions of time. Propofol infusion rates (in µg/kg/min) are shown underneath each plot. Note that in participants L514 and L585, bolus injections of propofol (doses in mg shown in italics; injection times indicated by arrows) were given per clinical considerations. Filled rectangles represent resting state data collection blocks; colors denote recording epochs corresponding to the three arousal states (red: awake; green: sedated; blue: unresponsive; gray: unassigned).

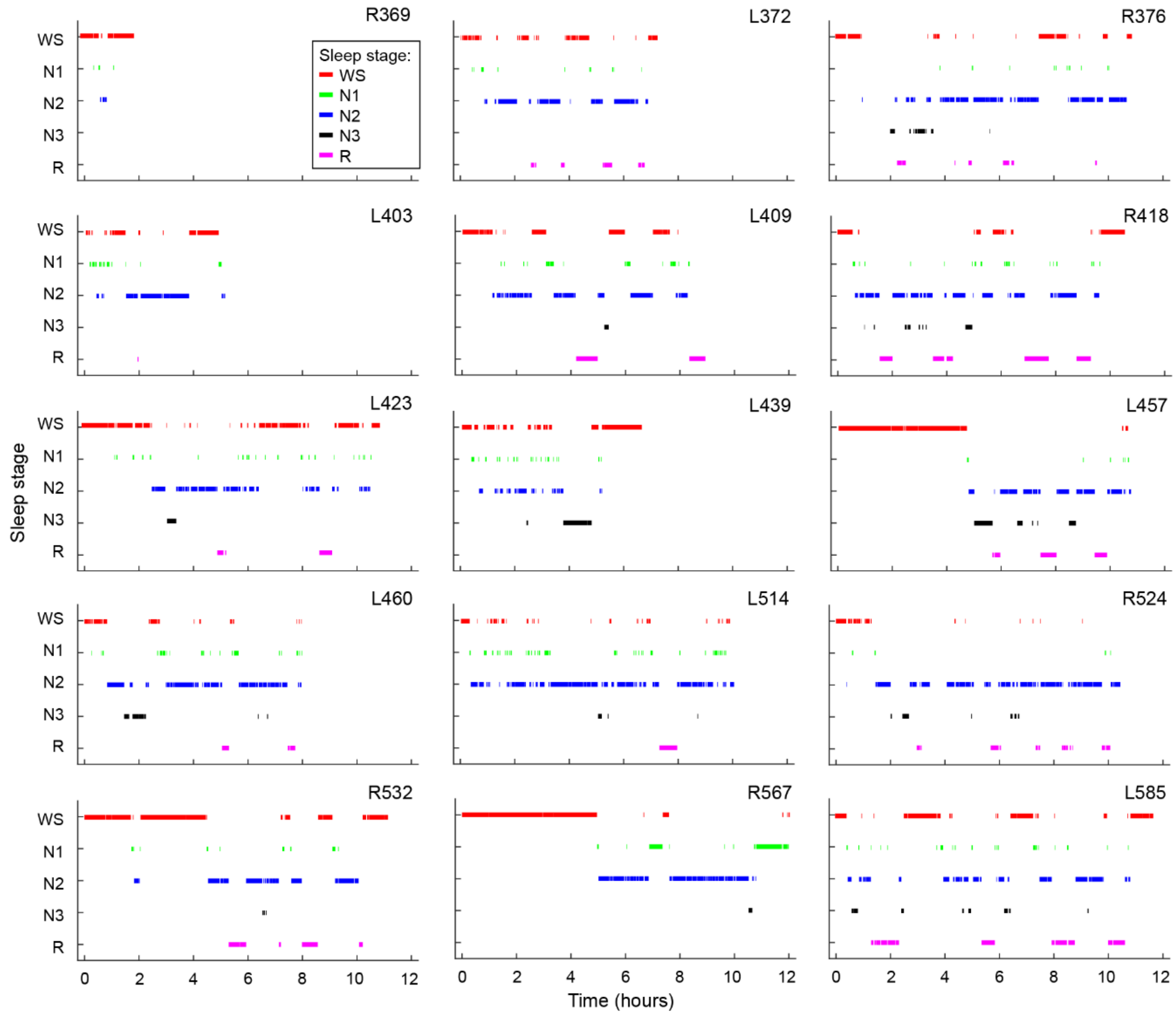


**Supplementary Figure 2. Sleep hypnograms in all participants.** Sleep staging was performed using standard polysomnography. Only shown are data segments used in the analysis, i.e., segments that were 60-seconds of continuous data with the same stage label, and that survived artifact rejection.

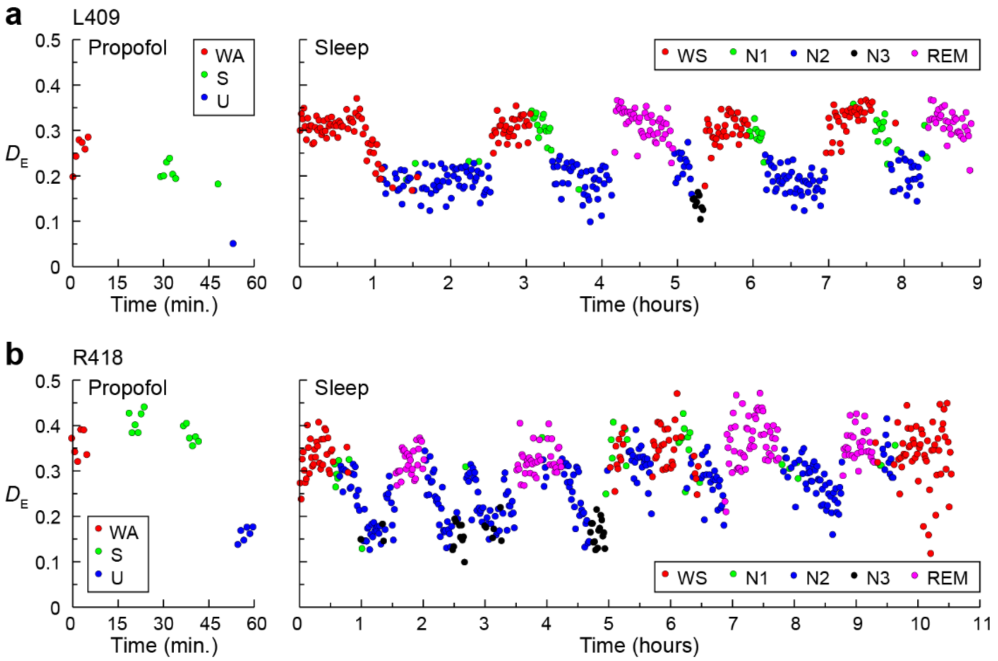


**Supplementary Figure 4. Time series of $D_E$ from two additional example participants.** Changes during anesthesia and sleep are shown in left and right panels, respectively. Each data point represents one minute of data. Exemplar data from participant L409 (**a**) and R418 (**b**).

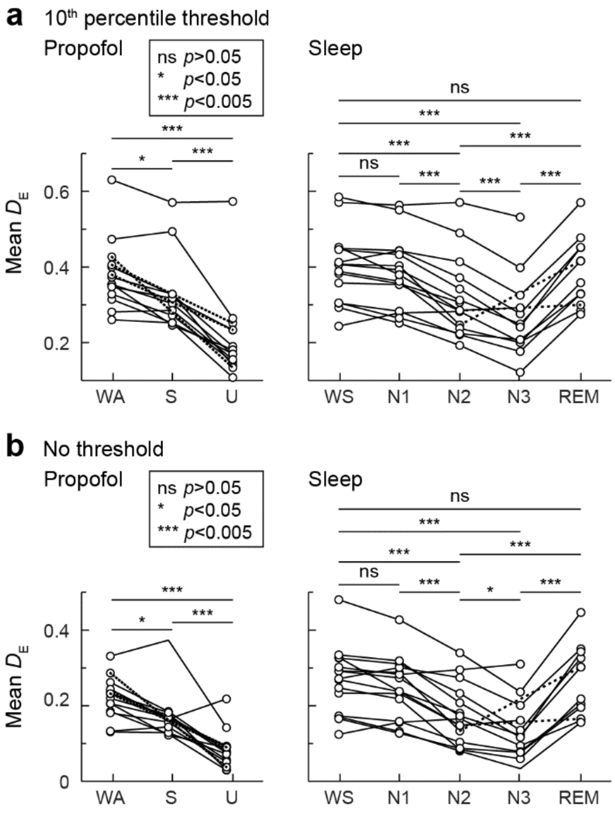


**Supplementary Figure 5. Changes in mean effective dimensionality are robust to the choice of threshold.** Conventions are the same as Figure 3. **a**: Using a more stringent threshold, keeping only the 10th percentile of connections in each row. **b**: Using a more permissive threshold where negative envelope correlations are set to zero but all positive correlations are retained.

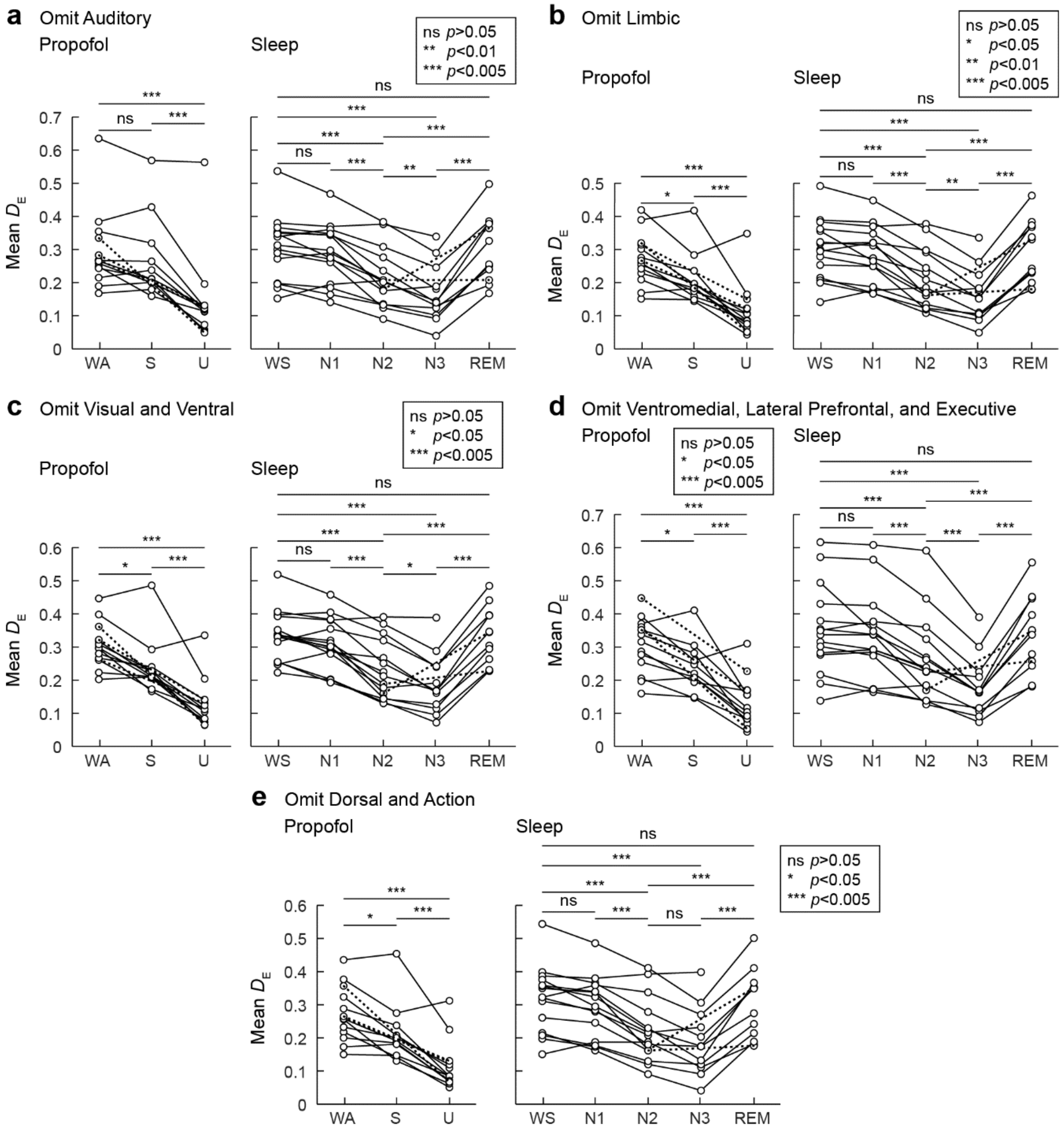


**Supplementary Figure 6. Changes in effective dimensionality are robust to exclusion of ROI groups.** Each panel exhibits a sensitivity analysis where all nodes from a given ROI group are excluded (**a**: Auditory; **b**: Visual; **c**: Dorsal; **d**: Ventral; **e**: Limbic; **f**: Executive; **g**: Action). The results are not qualitatively changed for any exclusion, indicating that no particularly influential ROI group is responsible for the overall changes observed. Conventions are the same as in Figure 3.

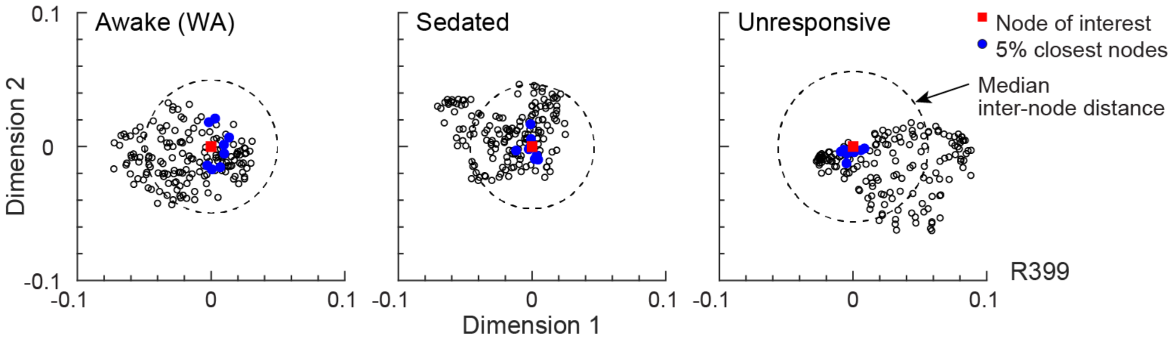


**Supplementary Figure 7. Normalized local distance.** We define “local distance” as the averaged distance between a node of interest and the 5% closest nodes, divided by the median distance to all nodes. This figure depicts this measure for one node for one subject, simplified to two dimensions for display. Axes reflect distance along the first two dimensions from the reference node (red square). The closest 5% nodes are marked (filled blue circles) as well as all other nodes (open circles). The closest nodes in the two dimensions shown are not necessarily the closest nodes in the full space. The radius of the dashed circle depicts the median Euclidean distance from reference node to other nodes. Compared to awake (WA), in Unresponsive the closest nodes are even closer, while the median is further.
